# Epilepsy gene therapy using non-integrating lentiviral delivery of an engineered potassium channel gene

**DOI:** 10.1101/298588

**Authors:** Albert Snowball, Elodie Chabrol, Robert C. Wykes, Andreas Lieb, Kevan S. Hashemi, Dimitri M. Kullmann, Matthew C. Walker, Stephanie Schorge

**Author notes:** These authors contributed equally (Albert Snowball, Elodie Chabrol, Robert C. Wykes). Joint corresponding authors.

## Abstract

Refractory focal neocortical epilepsy is a devastating disease for which there is frequently no effective treatment. Gene therapy represents a promising alternative, but treating epilepsy in this way involves irreversible changes to brain tissue, so vector design must be carefully optimized to guarantee safety without compromising efficacy. We set out to develop an epilepsy gene therapy vector optimized for clinical translation. The gene encoding the voltage-gated potassium channel Kv1.1, *KCNA1*, was codon-optimized for human expression and mutated to accelerate the channels’ recovery from inactivation. For improved safety, this engineered potassium channel (EKC) gene was packaged into a non-integrating lentiviral vector under the control of a cell type-specific *CAMK2A* promoter. In a blinded, randomized, placebo-controlled pre-clinical trial, the EKC lentivector robustly reduced seizure frequency in a rat model of focal neocortical epilepsy characterized by discrete spontaneous seizures. This demonstration of efficacy in a clinically relevant setting, combined with the improved safety conferred by cell type-specific expression and integration-deficient delivery, identify EKC gene therapy as ready for clinical translation in the treatment of refractory focal epilepsy.

## Introduction

Epilepsy affects over 60 million people worldwide (Ngugi et al., 2010). Even with optimal treatment approximately 30% remain resistant to pharmacotherapy (Kwan et al., 2011; Picot et al., 2008). The development of new anti-epileptic drugs in the last 20 years has had little impact on refractory epilepsy; people with inadequately controlled seizures continue to experience major co-morbidities, social exclusion, and an annual rate of sudden unexpected death in epilepsy (SUDEP) of 0.5-1% (Devinsky, 2011; Hoppe and Elger, 2011). Although surgical resection of the epileptogenic zone can result in seizure freedom, it is unsuitable for over 90% of refractory epilepsy patients (Lhatoo et al., 2003). Surgical intervention in focal neocortical epilepsy (FNE) is further complicated by the high risk of damage to eloquent regions of the cortex involved in functions such as memory, language, vision or fine motor control (Schuele and Lüders, 2008). People with FNE are therefore often left with very few, usually palliative, treatment options, and there is an urgent need to develop alternative therapies.

Gene therapy is one promising option (Kullmann et al., 2014), but major hurdles remain in achieving stable, predictable and safe transgene expression with viral vectors. Because focal seizures often arise from brain areas very close to eloquent cortex, lentiviral vectors, which generally lead to rapid, stable and, most importantly, spatially-restricted transgene expression (Lundberg et al., 2008), are an attractive delivery tool. In addition, the large packaging capacity of lentivectors allows a wide choice of promoter-transgene combinations (Kantor et al., 2014), which can further increase the specificity of expression. Hitherto, clinical trials with lentivectors for CNS disorders have been mainly restricted to *ex-vivo* treatment of hematopoietic stem cells (Biffi et al., 2013, 2013; Cartier et al., 2009). However, a recent trial using a lentivector injected directly into the striatum has demonstrated safety and tolerability in Parkinson’s disease, with evidence of decreased L-DOPA requirement (Palfi et al., 2014).

Early studies of gene therapy for epilepsy focused on acutely precipitated seizures, which often translate poorly (Galanopoulou et al., 2012). More recent strategies, mainly involving delivery of adeno-associated viral (AAV) vectors to models of temporal lobe epilepsy, have shown that the development of seizures after an epileptogenic insult (epileptogenesis) can be attenuated (Bovolenta et al., 2010; Haberman et al., 2003; Kanter-Schlifke et al., 2007; Lin et al., 2006; McCown, 2006; Nikitidou et al., 2014; Noé et al., 2008; Richichi et al., 2004; Woldbye et al., 2010). We have recently reported several approaches to gene therapy in a model of *epilepsia partialis continua* (EPC) induced by tetanus neurotoxin (TeNT) injection into the rat motor cortex (Kätzel et al., 2014; Wykes et al., 2012). In this model pathological high-frequency electrocorticographic (ECoG) activity is prominent, but discrete seizures lasting over 20 seconds are rare. Lentiviral overexpression of the human potassium channel Kv1.1, encoded by *KCNA1,* was highly effective at reducing pathological high frequency activity (Wykes et al., 2012). *In vitro* studies showed that Kv1.1 overexpression reduced both intrinsic neuronal excitability and glutamate release from transduced pyramidal neurons (Heeroma et al., 2009; Wykes et al., 2012). Importantly, both effects were graded, with neither neuronal excitability nor neurotransmitter release completely abolished. However, it remains unclear whether these graded effects on excitability and transmitter release, and the reduction of pathological ECoG activity in the motor cortex, can translate to the therapeutic suppression of intermittent discrete seizures.

Gene therapy based on overexpression of Kv1.1, or other proteins that reduce neuronal activity, requires effective targeting of transgene expression to excitatory neurons. Our previous work relied on driving *KCNA1* overexpression with a strong viral promoter, CMV. Although this promoter may bias expression to excitatory neurons in rat, recent data suggests it is not capable of doing so in non-human primates (Lerchner et al., 2014; Yaguchi et al., 2013). Restricting transgene expression to particular neuronal subtypes can however be achieved with the use of cell type specific promoters, but it is not yet known if these can support a level of expression sufficient to dampen neuronal activity. Finally, current clinical guidance seeks to reduce the risk of insertional mutagenesis associated with viral integration into the genome (Baum et al., 2004; Hacein-Bey-Abina et al., 2003), but there are few data indicating whether non-integrating lentiviral constructs can yield strong, stable transgene expression within the CNS.

To bring potassium channel gene therapy closer to the clinic, we have developed an optimized lentiviral vector designed to boost Kv1.1 expression and reduce its inactivation with an engineered *KCNA1* gene (EKC), and improve safety with a cell type specific (*CAMK2A*) promoter and non-integrating delivery vector (Rahim et al., 2009; Yáñez-Muñoz et al., 2006). The lentivector was tested for efficacy in a rat model of FNE characterised by long-lasting, discrete occipital cortex seizures (Chang et al., 2018). In a blinded, randomized, placebo-controlled pre-clinical trial, EKC gene therapy rapidly and persistently suppressed spontaneous seizures relative to a control lentivector without EKC.

## Results

### A pilot study shows that *KCNA1* gene therapy suppresses spontaneous seizures in a visual cortex epilepsy model

We first asked whether the CMV-driven *KCNA1* lentivector (CMV-*KCNA1*) used previously in a model of EPC (Wykes et al., 2012) was also effective in an epilepsy model characterized by discrete seizures. Epilepsy (Fig. 1A; Supp. Fig. 1) was induced in adult rats with a single injection of TeNT into the primary visual cortex. Seizures in this model typically last between 50 and 200 s, are accompanied by unilateral, bilateral or generalized convulsions, and evolve over several weeks before fading (Chang et al., 2018). To monitor local electrographic activity, a wireless ECoG transmitter was implanted with a subdural intracranial recording electrode positioned above the injection site. Two weeks after TeNT administration, following the establishment of epilepsy, animals were randomized into two groups and injected via a pre-implanted cannula with either the CMV-*KCNA1* lentivector or a control vector expressing only green fluorescent protein (GFP). Injections were delivered directly into the seizure focus and followed by a further 4 weeks of ECoG recording (Fig. 1B).

**Figure 1:**
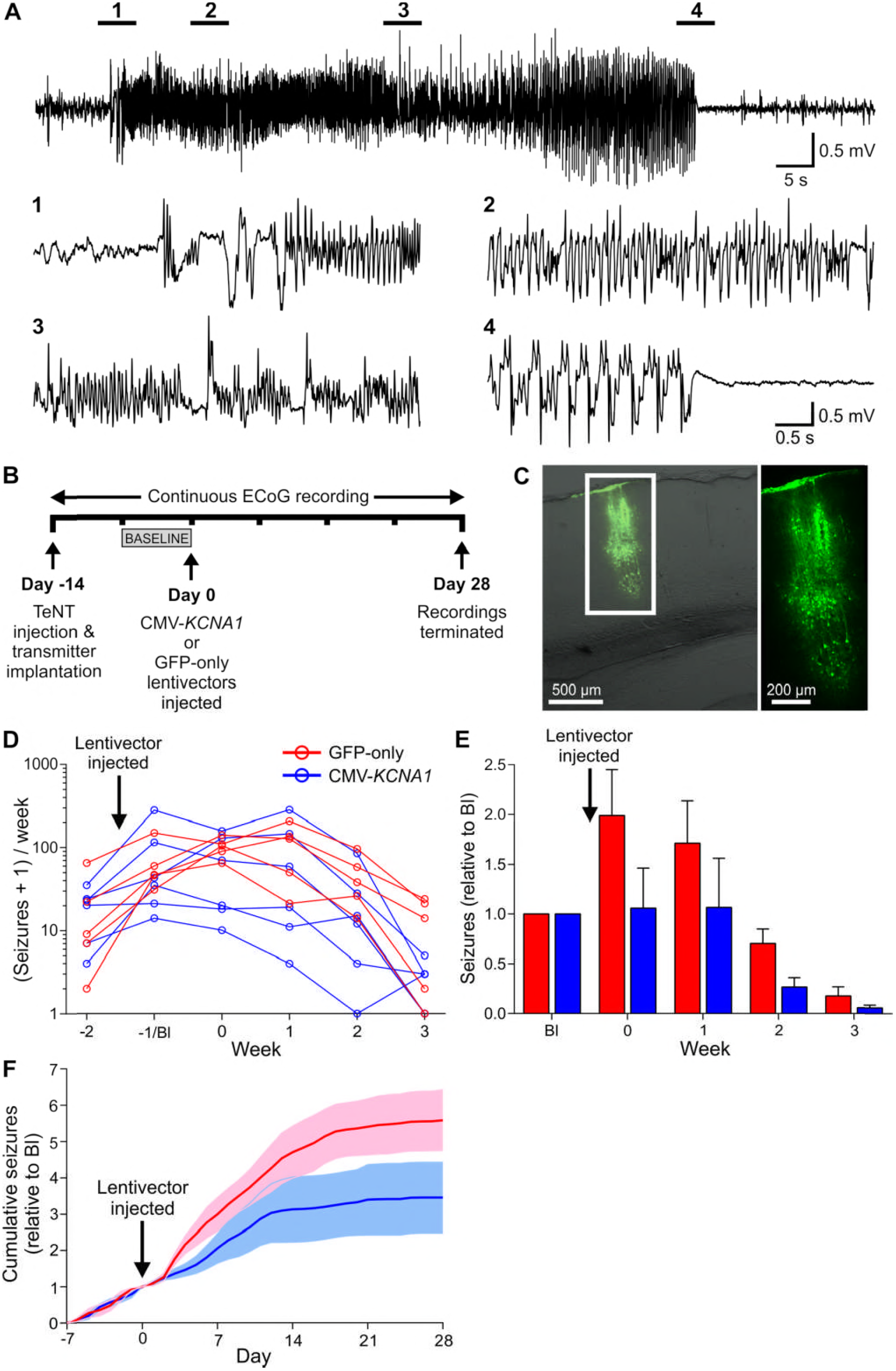
A pilot study suggests *KCNA1* gene therapy can suppress genuine discrete seizures in the visual cortex TeNT model of FNE. A. Representative occipital cortex seizure experienced by an adult rat 2 weeks after injection of TeNT into the primary visual cortex. Expanded sections are taken at the times indicated. Video footage of the behaviours associated with this electrographic seizure is provided in Fig EV1. B. Timeline highlighting key experimental milestones. C. Neuronal transduction with the CMV-*KCNA1* lentivector was restricted to a narrow column of cortex surrounding the site of injection additional images in Fig EV2. D. Number of seizures (per week) experienced by animals injected with the CMV-*KCNA1* lentivector (blue; n = 6) or its GFP-only control (red; n = 5). Data are plotted on a logarithmic scale after incrementing each seizure count by 1 to avoid zero values. E. Normalized seizure frequency (per week) for the two groups. The numbers of seizures experienced each week were normalized to the number experienced by each animal in the week preceding treatment (week Bl). F. Normalized cumulative seizure frequency (per day). Cumulative seizure counts were also normalized to the total number experienced in week Bl. Data in panels E and F are presented as mean ± the standard error of the mean (SEM).

The CMV-*KCNA1* lentivector transduced neurons within a narrow column of the cortex (Fig. 1C). As is typical of this model (Chang et al., 2018), the total number of seizures experienced by each animal over the 6 weeks of recording was highly variable (Fig. 1D). Consequently, to compare seizure frequency between the two treatment groups the numbers of seizures experienced each week were normalized to the number experienced in the week preceding treatment (week −1, or baseline (Bl) week). Despite the small sample size (6 treated vs. 5 controls), the CMV-*KCNA1* lentivector significantly reduced normalized seizure frequency compared to controls in the weeks following treatment (generalized log-linear mixed model on weeks 0 – 3, treatment*week interaction effect: F(1,40) = 4.851, p = 0.033; Fig. 1E). The therapeutic effect emerged rapidly; plots of normalized cumulative daily seizure frequency for the two groups diverged within 3 days of lentivector injection, consistent with rapid transgene expression, as seen previously in the motor cortex model (Fig. 1F).

This pilot study strongly suggests that *KCNA1* gene therapy can suppress spontaneous discrete seizures. However, the CMV-*KCNA1* lentivector tested is poorly suited for clinical translation. We therefore set out to develop an optimized vector with improved safety and efficacy.

### Design and characterization of an EKC gene therapy optimized for clinical translation

The transfer plasmid used to synthesize the optimized lentivector differed from the original CMV-*KCNA1* construct in several ways (Fig. 2A). The non-cell type specific CMV promoter was replaced with a 1.3 kb human *CAMK2A* promoter to bias expression to excitatory neurons (Dittgen et al., 2004; Yaguchi et al., 2013). The *KCNA1* gene was codon-optimized for expression in human cells, and mutated to introduce an I400V amino acid substitution normally generated by RNA editing. This substitution elicits a 20-fold increase in the rate at which Kv1.1 channels recover from inactivation (Bhalla et al., 2004). For pre-clinical evaluation, the coding sequence of a short-lived dscGFP reporter was linked to the EKC gene by a T2A element, which permits dual peptide expression from a single promoter. To ensure that the EKC construct could produce functional Kv1.1 channels, we performed whole-cell patch clamp recordings in transfected Neuro-2a cells, a line selected for its high *Camk2a* promoter activity. Robust non-inactivating Kv1.1 currents were recorded in cells transfected with the EKC plasmid (Fig. 2B).

**Figure 2:**
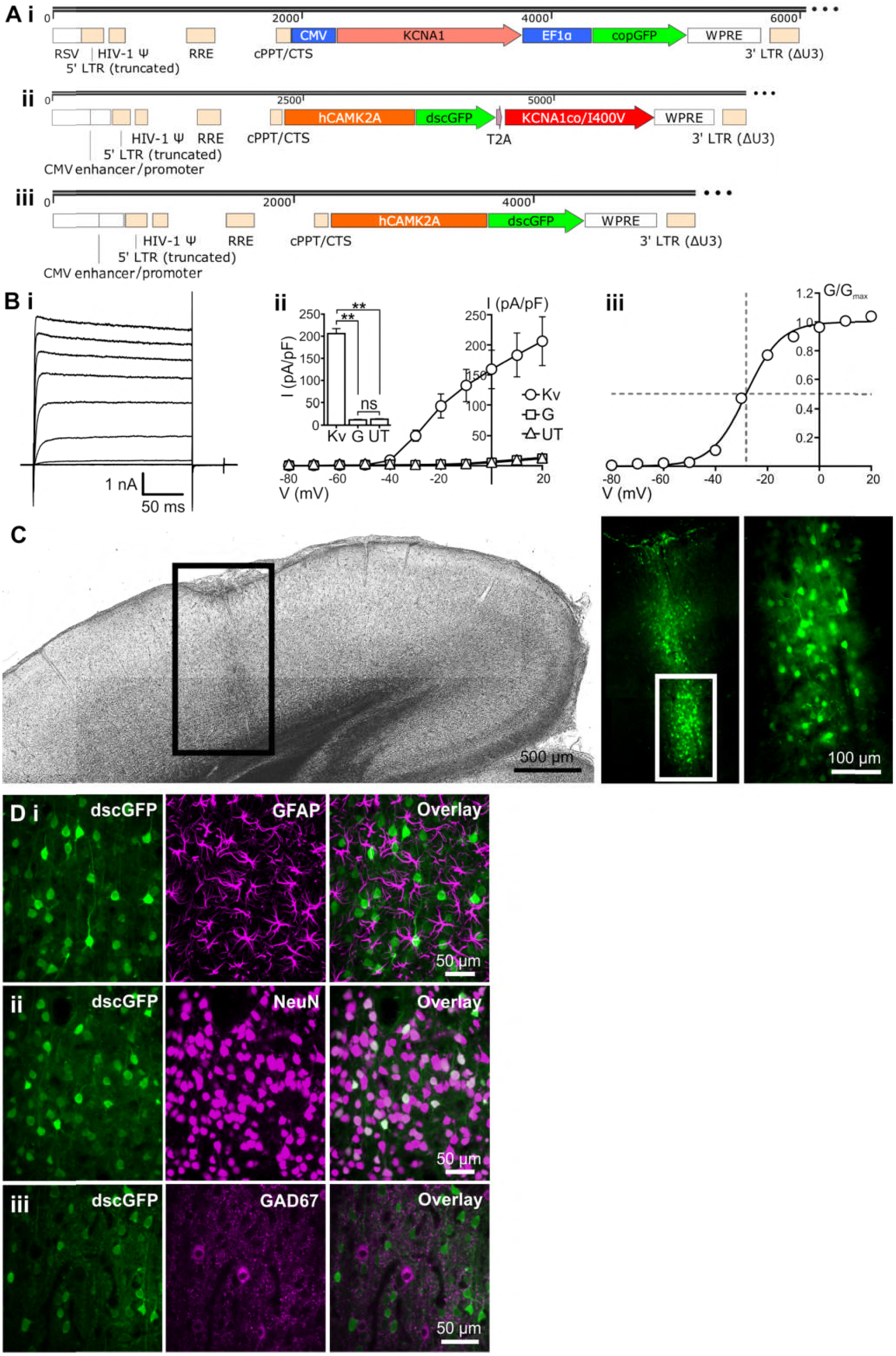
Design and characterization of an EKC gene therapy optimized for clinical translation. A. Lentiviral transfer plasmid maps for the CMV-*KCNA1* pilot vector (i), the optimized EKC vector (ii) and its dscGFP-only control (iii). Abbreviations: RSV – Rous sarcoma virus promoter; LTR – long terminal repeat; HIV-1 ll. – HIV-1 packaging signal; RRE – Rev response element; cPPT/CTS – central polypurine tract and central termination sequence; EF1a – elongation factor 1 a promoter; WPRE – woodchuck hepatitis virus post-transcriptional regulatory element. B. Heterologous expression of functional Kv1.1 channels from the optimized EKC transfer plasmid. (i): Representative current-time trace from a Neuro-2a cell transfected with the EKC transfer plasmid. (ii): Plot of mean current density against voltage for cells transfected with the EKC transfer plasmid (Kv; n=13), cells transfected with the dscGFP-only control plasmid (G; n=8), and untransfected controls (UT; n=10). Inset: histogram showing differences in current density between the three groups during the voltage step to +20 mV (Kv vs. UT: p=0.0013; Kv vs. G: p=0.0012; UT vs. G: p=0.82; ns = not significant; Welch’s one-way ANOVA with Games-Howell post-hoc tests). (iii): Plot of mean normalized conductance against voltage for cells transfected with the EKC transfer plasmid. Data are fit with a single Boltzmann function. The V_0.5_ (voltage of half-maximal conductance) of −28.2 mV is similar to values obtained from human embryonic kidney 293 (HEK293) cells transfected with CMV-driven, wild-type *KCNA1* (−32.8 ± 0.9 mV)(Tomlinson et al., 2013). All error bars represent SEM. C. Bright-field and fluorescence images of a brain slice from a rat injected in the left visual cortex with 1.25 μl (~3.0 × 10^6^ infectious units (IU)) of the EKC lentivector. The pattern of transduction is similar to that observed with the CMV-*KCNA1* vector. D. Immunohistochemical assessment of the cell type specificity of EKC expression. (i): There was no overlap between transduced neurons expressing dscGFP and astrocytes stained for GFAP. (ii): There was 100% overlap between dscGFP+ cells and neurons stained for NeuN. (iii): Minimal overlap was observed between dscGFP+ cells and inhibitory interneurons stained for GAD67.

The EKC transfer plasmid was packaged into a non-integrating lentiviral vector (Yáñez-Muñoz et al., 2006). When injected into the rat visual cortex, this lentivector drove strong, localized expression of the dscGFP reporter (Fig. 2C). Imaging of sequential brain slices yielded an estimated transduction volume of approximately 0.074 mm^3^ (Supp. Fig. 2). Immunohistochemistry revealed no visible overlap between dscGFP expression and glial fibrillary acidic protein (GFAP) staining (0/512 dscGFP+ cells stained for GFAP, n = 3 animals; Fig. 2Di). In contrast, all dscGFP+ cells stained positively for the neuronal marker NeuN (714/714, n = 3 animals; Fig. 2Dii). These data indicate that transgene expression from the EKC lentivector is restricted to neurons. There was minimal overlap between dscGFP expression and staining for glutamic acid decarboxylase 67 (GAD67), an enzymatic marker for GABAergic neurons (3/603 dscGFP+ cells stained for GAD67, n = 3 animals; Fig. 2Diii). This suggests that EKC transgene expression is largely restricted to excitatory neurons.

### EKC gene therapy reduces seizure frequency in a blinded, randomized pre-clinical trial

To test the therapeutic efficacy of the EKC lentivector, we designed a blinded, randomized, placebo-controlled pre-clinical trial, and selected normalized seizure frequency as the primary outcome measure. Eleven days after injection of TeNT into the visual cortex, 26 rats were randomized into two groups and injected via a pre-implanted cannula with either the EKC lentivector or its dscGFP-only control. ECoG recordings were continued for a further 4 weeks. The timeline was altered from that of the pilot study to treat after 11 days in order to capture the period when seizure activity is at its highest (2 – 4 weeks following TeNT injection) (Fig. 3A).

**Figure 3:**
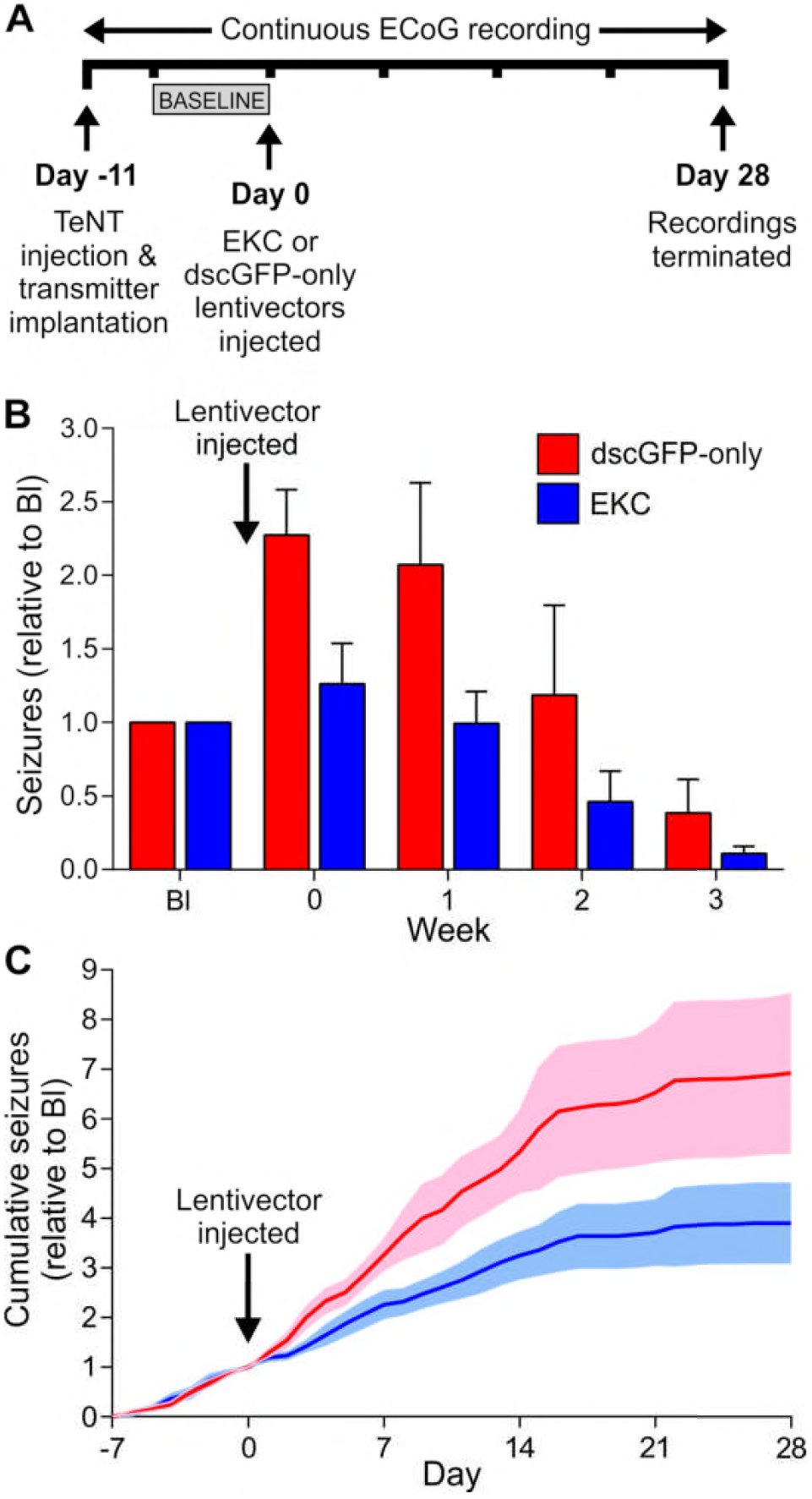
EKC gene therapy robustly reduces seizure frequency in a blinded, randomized, placebo-controlled pre-clinical trial. A. Timeline highlighting key experimental milestones. Note the injection of lentiviral vectors 11 days after TeNT delivery. B. Normalized seizure frequency (per week) for animals treated with the EKC lentivector (blue; n = 7/6) or its dscGFP-only control (red; n = 11). C. Normalized cumulative seizure frequency (per day). Data in panels B and C are presented as mean± SEM.

To minimize the confounding influence of animals that displayed a very low seizure frequency prior to treatment, subjects were excluded if they exhibited fewer than five seizures in the week preceding lentiviral delivery (the baseline week). This criterion, applied before unblinding, led to the exclusion of eight animals (6 EKC, 2 control). Of the remaining 18, all but one survived for the duration of recording. This rat (from the EKC group) was culled in the final week due to detachment of its headpiece. However, because the subject had already passed through the period of peak seizure activity, and in order to maximise the amount of data obtained from the study, this incomplete dataset was included in the overall analysis. Again, this decision was made before unblinding.

There was no significant difference between the treatment groups in the number of seizures experienced in the week preceding virus injection (control median = 11 (IQR 10 – 26), EKC median = 10 (IQR 7.5 – 12); Mann Whitney U test, p = 0.185). Analysis of the primary outcome measure indicated that EKC therapy robustly decreased normalized seizure frequency compared to controls in the weeks following treatment (generalized log-linear mixed model on weeks 0 – 3, treatment*week interaction effect: F(1,67) = 29.704, p < 0.001; Fig. 3B). The size of the effect was larger than that observed in the pilot study, suggesting that the EKC gene is more effective than its wild-type *KCNA1* counterpart at suppressing neuronal hyperexcitability. As in the pilot study, the reduction in seizure frequency lasted for the duration of recording, and the absolute effect size only decreased as seizures abated in the control group. Again the therapeutic effect emerged rapidly, with plots of normalized cumulative daily seizure frequency for the two groups diverging 2 days after treatment (Fig. 3C).

## Discussion

EKC gene therapy represents an effective new treatment for focal neocortical seizures in a format adapted to improve safety and suitability for human use. The results presented here provide strong justification for the further clinical development of EKC therapy.

We have previously shown that overexpression of Kv1.1 can reduce the frequency of brief (< 1 s), high-frequency epileptiform discharges in a motor cortex TeNT model of EPC (Wykes et al., 2012). However, this study did not investigate whether Kv1.1 overexpression could inhibit discrete seizures lasting 1 – 2 minutes, more typical of common forms of focal epilepsy. We show here, in two independent trials, that Kv1.1 overexpression is indeed sufficient to reduce the frequency of seizures, although interestingly seizure duration was not altered. This is most simply explained by proposing that seizure initiation is rapidly accompanied by propagation to other brain areas, beyond the lentivector-treated region, consistent with the convulsions seen in the majority of seizures in the visual cortex TeNT model (Chang et al., 2018).

Injection of TeNT into the occipital cortex induced seizures that lasted markedly longer (50 – 150 s) than the epileptiform bursts evoked by TeNT injection in motor cortex (< 1 s) (Wykes et al., 2012). The difference, which parallels that seen with occipital lobe seizures and EPC in human patients, may be a consequence of different connectivity in the occipital and motor cortices. Further studies will be needed to determine how cortical architecture impacts the type of epileptiform activity induced by TeNT insult.

Lentiviral gene therapy approaches are becoming more common in CNS disorders, and have shown good safety and tolerability even in extended trials (Palfi et al., 2014). However, a potential safety concern with retroviral vectors is the inherent risk of insertional mutagenesis (Baum et al., 2004; Hacein-Bey-Abina et al., 2003). This risk can be minimized by rendering vectors integration-deficient. The popularity of non-integrating lentiviruses for therapeutic gene transfer is growing, and the vectors have already demonstrated pre-clinical efficacy in the treatment of degenerative retinal disease and haemophilia B (Suwanmanee et al., 2014; Yáñez-Muñoz et al., 2006). The non-integrating EKC lentivirus described here drove strong, localized transgene expression after direct injection into the rat neocortex, and rapidly and persistently suppressed focal seizure activity. This supports the use of integration-deficient vectors as safe, effective delivery tools for gene therapy of neurological disease.

In the case of epilepsy, an additional safety concern is the possibility of potassium channel overexpression in interneurons, which could aggravate seizure activity by exacerbating rather than attenuating local excitability. To mitigate this risk we have used a human *CAMK2A* promoter that in rats led to very little expression in GABAergic cells. Promoter specificity can differ between species (Lerchner et al., 2014; Yaguchi et al., 2013), and the specificity of the human *CAMK2A* promoter for excitatory glutamatergic neurons will ultimately need to be validated in the human brain. Evidently, if EKC gene therapy is to progress to the clinic, such validation will need to be performed in the absence of a fluorescent reporter.

Because the role of potassium channels, including Kv1.1, in regulating neuronal excitability is conserved across a broad range of neurons, potassium channel overexpression may hold therapeutic promise in the treatment of other diseases characterized by neuronal hyperexcitability. There is currently an unmet clinical need for new treatments of chronic pain, and a variety of gene therapy approaches aimed at reducing the excitability of dorsal root ganglion neurons have already demonstrated pre-clinical efficacy (Snowball and Schorge, 2015). Other disorders such as Parkinson’s disease are associated with excessive activity in specific groups of neurons (Lobb, 2014), and could be candidates for treatment with an appropriate combination of potassium channel subtype and cell type specific promoter.

## Materials and methods

### Molecular biology

Lentiviral transfer plasmids were constructed using standard subcloning techniques. *KCNA1* was codon optimized for human expression using GeneOptimizer® software, and synthesized using GeneArt^®^ (Thermo Fisher Scientific). All plasmids were fully sequenced before use. Sequences are available on request.

### Voltage clamp recordings

Neuro-2a cells were grown in Gibco^®^ Dulbecco’s Modified Eagle Medium (DMEM) + GlutaMAX™ (Thermo Fisher Scientific) supplemented with 10% heat-inactivated foetal bovine serum (Thermo Fisher Scientific), 1% penicillin/streptomycin (Thermo Fisher Scientific) and 1% non-essential amino acids (Sigma). Cultures were maintained in logarithmic growth phase in a humidified 5% CO_2_ atmosphere at 37 °C. Transfections were performed according to the manufacturer’s instructions using TurboFect™ transfection reagent (Thermo Fisher Scientific). Transfected cells were plated onto 13 mm borosilicate glass coverslips (VWR). Coverslips were placed into the chamber of a BX51WI fixed-stage upright microscope equipped with UMPLFLN 10× and LUMPLFLN 40× water-immersion objectives (Olympus). Coverslips were submerged in a static bath of extracellular solution with the following composition (in mM): 140 NaCl, 4 KCl, 1.8 CaCl_2_,2 MgCl_2_, 10 HEPES (pH 7.35, osmolarity ~301 mOsm/L). Filamented borosilicate glass micropipettes (GC150-F; Warner Instruments) were pulled to tip resistances between 2.0 and 3.0 MΩ using a P-97 Flaming/Brown micropipette puller (Sutter Instrument Company). Micropipettes were filled with an intracellular solution of the following composition (in mM): 140 KCl, 10 HEPES, 10 EGTA (pH 7.35, osmolarity ~291 mOsm/L). Macroscopic currents were recorded under voltage clamp using the whole-cell patch clamp configuration. The voltage step protocol used was as follows: cells were held at a resting potential of −80 mV and currents evoked by 200 ms depolarising steps delivered in 10 mV increments up to +20 mV. A 40 ms hyperpolarising step to −100 mV was included before returning to baseline. Data were filtered at 3 kHz and acquired at 10 kHz using WinWCP software (J. Dempster, University of Strathclyde) and an Axon Multiclamp 700B amplifier (Molecular Devices). Series resistance compensation was employed throughout, with prediction and correction components adjusted to 80% and the bandwidth set to 1.2 kHz. Cells with series resistance greater than 10 MΩ were excluded from the analysis. All recordings were made at room temperature (23 – 26 °C). The liquid junction potential, calculated to be +4.1 mV, was left uncorrected. Leak currents were minimal and left unsubtracted.

For analysis, evoked currents were taken as the steady-state current in the last 40 ms of each voltage step. Baseline holding currents were subtracted before division by cell capacitance to generate current density values. To calculate normalized conductance, the current density at each voltage step was divided by the step potential minus the potassium reversal potential (−91.34 mV). This generates raw conductance values that are corrected for the variation in K^+^ driving force which accompanies stepwise changes in membrane potential. Plots of raw conductance against voltage for each EKC-transfected cell were fit with individual Boltzmann functions given by the equation:

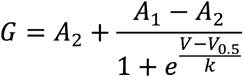

where G is the conductance, V the voltage, A_1_ the initial (minimum) conductance, A_2_ the final (maximum) conductance, V_0.5_ the voltage of half-maximal conductance, and k the slope factor. Raw conductance values were normalized to A_1_ and A_2_ of their own Boltzmann functions. Normalized conductance was then plotted against voltage for all EKC-transfected cells and mean values fit with a single Boltzmann function (Fig. 2Biii).

### Lentiviral synthesis

The CMV-*KCNA1* lentivector was identical to that used in Wykes *et al.*, 2012 (Wykes et al., 2012). For the EKC lentivector and its dscGFP-only control, HEK293T producer cells were grown in Gibco^®^ DMEM + GlutaMAX™ supplemented with 10% heat-inactivated foetal bovine serum and 1% penicillin/streptomycin. Cultures were maintained in logarithmic growth phase in a humidified 5% CO_2_ atmosphere at 37 °C. Cells were split every 3 – 4 days using 0.05% Trypsin-EDTA (Thermo Fisher Scientific) and never grown for more than 15 passages. Cells were co-transfected with pMDG-VSV.G, pCMVdR8.74^D64V^ and either the EKC transfer plasmid or its dscGFP-only control. The mass ratio of envelope to packaging to transfer plasmids was 1: 2.5: 1.5. Transfections were performed according to the manufacturer’s instructions using Lipofectamine^®^ 2000 (Thermo Fisher Scientific). The transfection medium was replaced after 18 hours. Two media harvests were collected, at 40 hours and 60 hours after transfection. Harvested media were pre-cleaned by centrifugation at 1000 rpm for 3 minutes at 4 °C and filtered through 0.45 μm micropores. Media were overlaid on a sucrose solution with the following composition (in mM): 50 Tris-HCl, 100 NaCl, 0.5 EDTA (pH 7.4, 10% w/v sucrose), and centrifuged at 20,000 rpm for 2 hours at 4 °C. Lentiviral pellets were resuspended in sterile PBS, aliquoted, snap-frozen and stored at −80 °C. Viral titre was approximated using the Lenti-X™ p24 rapid titer kit (Clontech). Each titration was performed in triplicate with 3 separate aliquots. Estimated titres were 2.42 × 10^9^ IU/ml (EKC lentivector) and 4.26 × 10^9^ IU/ml (dscGFP-only control).

### Surgical procedures

All experiments were performed in accordance with the United Kingdom Animals (Scientific Procedures) Act 1986. Adult male rats (Sprague Dawley; 300-400g) were anesthetized and placed into a stereotaxic frame (Kopf). 15 ng of TeNT was injected into layer 5 of the right visual cortex in a final volume of 1.0 µl at a rate of 100 nl/min (coordinates: 3 mm lateral, 7 mm posterior of bregma; 1.0 mm deep from the pia). An ECoG transmitter (A3028E; Open Source Instruments, MA, USA) was implanted subcutaneously with a subdural intracranial recording electrode positioned above the injection site. A reference electrode was implanted in the contralateral hemisphere. A cannula (Plastics One) was positioned above the injection site for delivery of lentiviral vectors 11 or 14 days later. Each rat received a maximum of 2.0 µl of lentivirus injected directly into the seizure focus. Animals injected with TeNT were housed separately in Faraday cages for the duration of the study.

### ECoG acquisition and analysis

ECoG was recorded continuously for up to 6 weeks after surgery. Data were acquired using A3028E implantable transmitters (0.3 – 160 Hz, 512 samples/s) and ancillary receivers and software (Open Source Instruments, Inc.). The method of seizure detection differed for the pilot study and the final pre-clinical trial. For the pilot study, ECoG traces were first divided into 1 s epochs. Four metrics (power, coastline, intermittency and coherence) were then quantified for each epoch, and their values compared to those from a user-curated library of epochs validated by video as representing seizure activity (Wykes et al., 2012). Matched values were fed into a consolidation script that returned all instances of 5 or more sequential epochs identified as containing seizure activity. All seizures in the consolidation output were verified by an experimenter. For the final pre-clinical trial, 6 metrics were quantified for each epoch (power, coastline, intermittency, coherence, asymmetry and rhythm) and all matched values were manually checked for seizure activity without the use of a consolidation script. Seizure counts in this trial were performed by an experimenter blinded to the treatment. For all datasets the minimum duration for a seizure was set at 10 s.

### Immunohistochemistry

One week after lentivirus injection rats were terminally anesthetized with sodium pentobarbital (Euthatal; Merial) and transcardially perfused with cold (4 °C) heparinized PBS (80 mg/L heparin sodium salt; Sigma) followed by 4% paraformaldehyde (PFA) in PBS (Santa Cruz Biotechnology). Brains were removed and post-fixed in 4% PFA at 4 °C for a further 24 hours. After washing in PBS brains were sliced into 70 μm coronal sections using a vibrating microtome (Leica) and stored free-floating at 4 °C in PBS plus 0.02% sodium azide (Sigma). For antibody staining, slices were permeabilized for 20 minutes in PBS plus 0.3% Triton X-100 (Sigma) before blocking for 1 hour in PBS plus 0.3% Triton X-100, 1% bovine serum albumin (Sigma) and 4% goat serum (Sigma). Slices were incubated overnight at 4 °C in PBS plus 0.3% Triton X-100 and a rabbit anti-NeuN (diluted 1:750; ab177487; Abcam), mouse anti-GFAP (diluted 1:500; MAB3402; Merck Millipore) or mouse anti-GAD67 (diluted 1:500; MAB5406; Merck Millipore) primary antibody. After three 10 minute washes in PBS, slices were incubated at room temperature for 3 hours in PBS plus the relevant Alexa Fluor^®^ 594-conjugated secondary antibody (goat anti-rabbit (A-11037; Thermo Fisher Scientific) or goat anti-mouse (A-11005; Thermo Fisher Scientific); both diluted 1:750). After a further three 10 minute washes in PBS, slices were mounted onto plain glass microscope slides (Thermo Fisher Scientific) using Vectashield^®^ HardSet™ mounting medium (Vector Laboratories) and borosilicate glass coverslips (VWR). Bright-field and fluorescence images were acquired using one of two microscopes: an Axio Imager A1 fluorescence microscope (Axiovision LE software) equipped with 2.5×, 10× and 40× EC Plan-Neofluar non-immersion objectives, or an inverted LSM 710 confocal laser scanning microscope (ZEN 2009 software) equipped with 40× and 63× EC Plan-Neofluar oil-immersion objectives (all Zeiss). For the confocal microscope, dscGFP and Alexa Fluor^®^ 594 were excited with the 488 nm and 561 nm lines of an argon or diode pumped solid state (DPSS) laser, respectively. All image processing was performed using ImageJ software. Composite images were assembled using the MosaicJ ImageJ plugin.

### Statistics

Data from the pilot study were used to determine sample sizes for the final pre-clinical trial. We estimated that the maximal weekly seizure frequency would double from baseline, and we wished to detect with 80% power a 40% reduction from this maximum at p < 0.05. Given a mean baseline weekly seizure frequency of 5 or above, a modification of Lehr’s formula (Lehr, 1992) for the Poisson distribution suggested 7 – 8 animals per group would be sufficient to detect a reduction in seizure frequency from 10 to 6 per week. Our modified Lehr’s formula is given by the following equation:

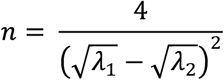

where n is the size of each sample (treatment group), λ_1_ the mean weekly seizure frequency before treatment, and λ_2_ the mean weekly seizure frequency after treatment.

Efficacy of treatment data (Fig. 1E, 3B) were analysed using a generalized log-linear mixed model with random effect of animal (autoregressive covariance) and fixed effects of treatment group, week, and the interaction between treatment group and week. Seizure counts in the week preceding treatment were compared using a Mann Whitney U test. Current densities at +20 mV (Fig. 2Bii) were compared using a Welch’s one-way ANOVA followed by Games-Howell post-hoc tests.

## Acknowledgments

We thank G. Schiavo (UCL Institute of Neurology) for the gift of TeNT, S. Hart (UCL Institute of Child Health) for the Neuro-2a cells, and A. J. Thrasher and W. Qasim (UCL Institute of Child Health) for the pMDG-VSV.G and pCMVdR8.74^D64V^ plasmids. We are grateful to J. Cornford for technological assistance in the visualisation of example seizures, and for the animal care provided by members of our Biological Services Unit. This work was supported by the Medical Research Council, the Wellcome Trust, Epilepsy Research UK, a Marie Sktodowska-Curie Actions Research Fellowship, and a Royal Society University Research Fellowship.

## Author contributions

AS designed, synthesized and characterized the EKC lentivector, and analyzed and interpreted data from the final pre-clinical trial. EC performed the final pre-clinical trial and analyzed and interpreted data from it. RCW performed the pilot study and analyzed and interpreted data from it. AL provided technical assistance for video/ECoG recordings. KSH designed and built the ECoG recording system, and analyzed and interpreted data from the pilot study. SS, DMK and MCW designed the study, supervised the experiments and interpreted the data. AS, SS, DMK and MCW wrote the manuscript with input from all co-authors.

## The Paper Explained

### Problem

Focal neocortical epilepsy is a serious and common disease that is frequently resistant to anti-epileptic drugs. Gene therapy is a promising treatment alternative to surgery to remove the seizure focus. In previous work, overexpression of *KCNA1*, encoding the voltage-gated potassium channel Kv1.1, suppressed pathological, high-frequency brain activity evoked by injecting tetanus toxin into the rat motor cortex. However, several features of the lentiviral vector used to deliver the *KCNA1* gene made it unsafe for human administration, and it was unclear if the reduction of pathological high-frequency activity would extend to a suppression of discrete seizures. We therefore developed a lentiviral vector optimized for clinical translation, and tested its effectiveness in a model of epilepsy characterized by discrete seizures.

### Results

To boost the efficacy of the gene therapy, *KCNA1* was codon-optimized for human expression and mutated to accelerate the recovery of Kv1.1 channels from inactivation. To improve safety, this engineered potassium channel gene was placed under the transcriptional control of a *CAMK2A* promoter to restrict expression to excitatory neurons, and packaged into a non-integrating lentiviral vector to reduce the risk of insertional mutagenesis. In a blinded, randomized, placebo-controlled pre-clinical trial, the EKC lentivector robustly reduced seizure frequency in a rat model of focal neocortical epilepsy characterized by discrete seizures.

### Impact

This demonstration of the anti-epileptic efficacy of *KCNA1* gene therapy in a clinically relevant setting, combined with the improved safety conferred by cell type specific expression and integration-deficient delivery, suggests EKC gene therapy is well placed for clinical translation in the treatment of drug-resistant focal epilepsy. To our knowledge this study represents the first successful use of a non-integrating lentiviral vector to treat an experimental model of a neurological disease, and addresses a major treatment gap for over 5 million people suffering from refractory focal epilepsy worldwide.

---

See separate .mp4 file for Expanded View Figure 1.

**Fig EV1: Time-locked video-ECoG recording of a spontaneous discrete seizure in the visual cortex TeNT model of FNE.**

As previously described (Chang et al., 2018) electrographic seizures in this model were accompanied by overt behaviours including sudden increases in arousal, repetitive eye blinking, rearing, bilateral limb twitching and wet dog shakes. Displayed is a representative video-ECoG recording of a rat experiencing a focal to bilateral tonic-clonic seizure.

**Fig EV2:**
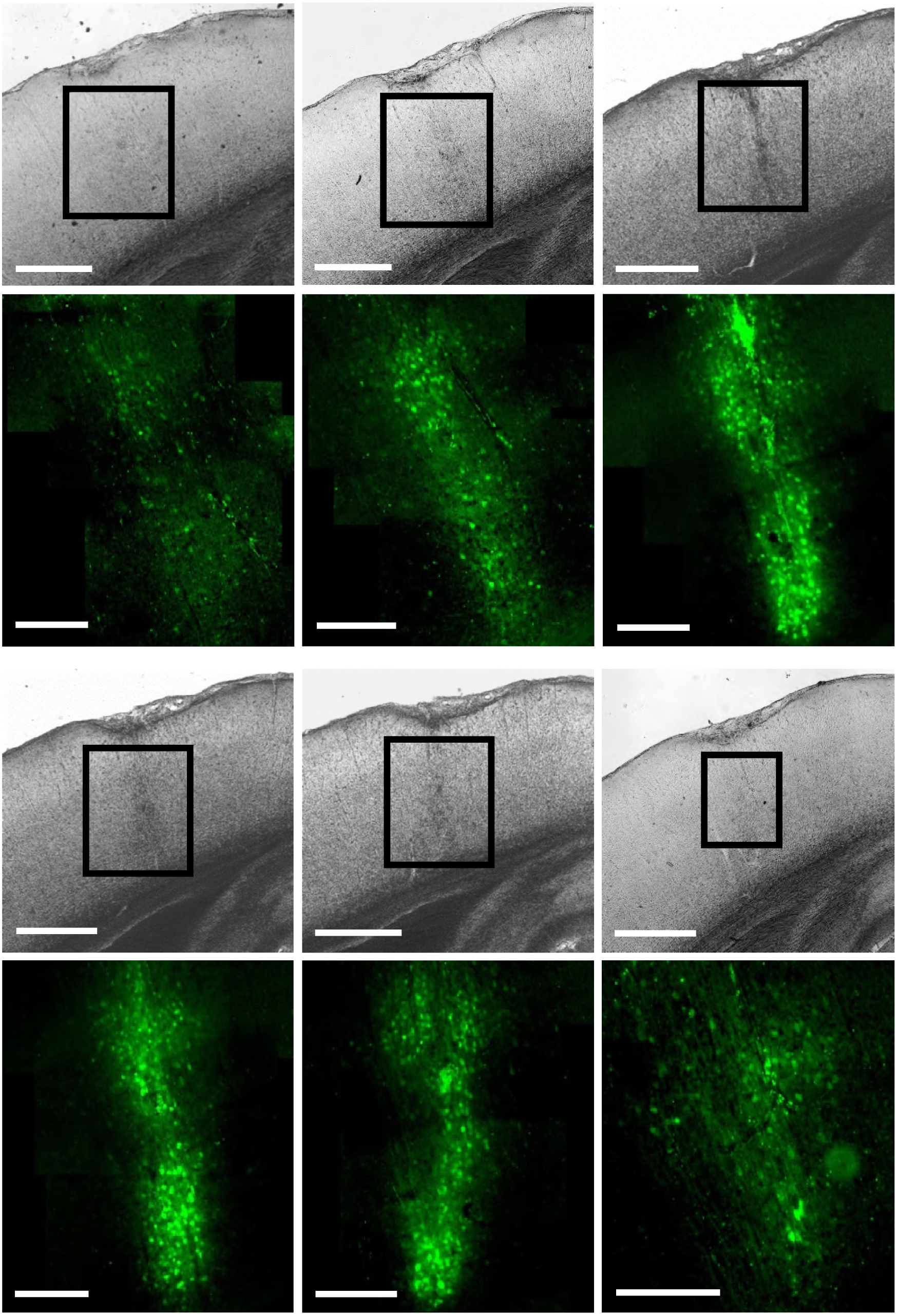
Spread of transduction with the EKC lentivector. Bright-field and fluorescence images of 6 sequential left-hemisphere visual cortex slices (70 μm thick) from a rat brain injected with 1.25 μl (~3.0 × 10^6^ IU) of the EKC lentivector. Slices are ordered from top left (rostral) to bottom right (caudal). Scale bars represent 600 μm and 200 μm for bright-field and fluorescence images, respectively.

## Bibliography

Baum, C., von Kalle, C., Staal, F.J.T., Li, Z., Fehse, B., Schmidt, M., Weerkamp, F., Karlsson, S., Wagemaker, G., and Williams, D.A. (2004). Chance or necessity? Insertional mutagenesis in gene therapy and its consequences. Mol. Ther. J. Am. Soc. Gene Ther. 9, 5–13.

Bhalla, T., Rosenthal, J.J.C., Holmgren, M., and Reenan, R. (2004). Control of human potassium channel inactivation by editing of a small mRNA hairpin. Nat. Struct. Mol. Biol. 11, 950–956.

Biffi, A., Montini, E., Lorioli, L., Cesani, M., Fumagalli, F., Plati, T., Baldoli, C., Martino, S., Calabria, A., Canale, S., et al. (2013). Lentiviral Hematopoietic Stem Cell Gene Therapy Benefits Metachromatic Leukodystrophy. Science 341, 1233158.

Bovolenta, R., Zucchini, S., Paradiso, B., Rodi, D., Merigo, F., Mora, G.N., Osculati, F., Berto, E., Marconi, P., Marzola, A., et al. (2010). Hippocampal FGF-2 and BDNF overexpression attenuates epileptogenesis-associated neuroinflammation and reduces spontaneous recurrent seizures. J. Neuroinflammation 7, 81.

Cartier, N., Hacein-Bey-Abina, S., Bartholomae, C.C., Veres, G., Schmidt, M., Kutschera, I., Vidaud, M., Abel, U., Dal-Cortivo, L., Caccavelli, L., et al. (2009). Hematopoietic stem cell gene therapy with a lentiviral vector in X-linked adrenoleukodystrophy. Science 326, 818–823.

Chang, B.-L., Leite, M., Snowball, A., Chabrol, E., Leib, A., Walker, M.C., Kullmann, D.M., Schorge, S., and Wykes, R.C. (2014). Semiology, clustering, periodicity and natural history of seizures in an experimental visual cortical epilepsy model. BioRxiv 289256.

Devinsky, O. (2011). Sudden, unexpected death in epilepsy. N. Engl. J. Med. 365, 1801–1811.

Dittgen, T., Nimmerjahn, A., Komai, S., Licznerski, P., Waters, J., Margrie, T.W., Helmchen, F., Denk, W., Brecht, M., and Osten, P. (2004). Lentivirus-based genetic manipulations of cortical neurons and their optical and electrophysiological monitoring in vivo. Proc. Natl. Acad. Sci. U. S. A. 101, 18206–18211.

Galanopoulou, A.S., Buckmaster, P.S., Staley, K.J., Moshé, S.L., Perucca, E., Engel, J., Jr, Löscher, W., Noebels, J.L., Pitkänen, A., Stables, J., et al. (2012). Identification of new epilepsy treatments: issues in preclinical methodology. Epilepsia 53, 571–582.

Haberman, R.P., Samulski, R.J., and McCown, T.J. (2003). Attenuation of seizures and neuronal death by adeno-associated80virus vector galanin expression and secretion. Nat. Med. 9, 1076–1080.

Hacein-Bey-Abina, S., Von Kalle, C., Schmidt, M., McCormack, M.P., Wulffraat, N., Leboulch, P., Lim, A., Osborne, C.S., Pawliuk, R., Morillon, E., et al. (2003). LMO-associated clonal T cell proliferation in two patients after gene therapy for SCID-X. Science 302, 415–419.

Heeroma, J.H., Henneberger, C., Rajakulendran, S., Hanna, M.G., Schorge, S., and Kullmann, D.M. (2009). Episodic ataxia type 1 mutations differentially affect neuronal excitability and transmitter release. Dis. Model. Mech. 2, 612–619.

Hoppe, C., and Elger, C.E. (2011). Depression in epilepsy: a critical review from a clinical perspective. Nat. Rev. Neurol. 7, 462–472.

Kanter-Schlifke, I., Georgievska, B., Kirik, D., and Kokaia, M. (2007). Seizure Suppression by GDNF Gene Therapy in Animal Models of Epilepsy. Mol. Ther. 15, 1106–1113.

Kantor, B., McCown, T., Leone, P., and Gray, S.J. (2014). Chapter Two - Clinical Applications Involving CNS Gene Transfer. In Advances in Genetics, J.C.D. and S.F.G. Theodore Friedmann, ed. (Academic Press), pp. 71–124.

Kätzel, D., Nicholson, E., Schorge, S., Walker, M.C., and Kullmann, D.M. (2014). Chemical-genetic attenuation of focal neocortical seizures. Nat. Commun. 5, 3847.

Kullmann, D.M., Schorge, S., Walker, M.C., and Wykes, R.C. (2014). Gene therapy in epilepsy-is it time for clinical trials? Nat. Rev. Neurol. 10, 300–304.

Kwan, P., Schachter, S.C., and Brodie, M.J. (2011). Drug-resistant epilepsy. N. Engl. J. Med. 365, 919–926.

Lehr, R. (1992). Sixteen S-squared over D-squared: A relation for crude sample size estimates. Stat. Med. 11, 1099–1102.

Lerchner, W., Corgiat, B., Der Minassian, V., Saunders, R.C., and Richmond, B.J. (2014). Injection parameters and virus dependent choice of promoters to improve neuron targeting in the nonhuman primate brain. Gene Ther.

Lhatoo, S.D., Solomon, J.K., McEvoy, A.W., Kitchen, N.D., Shorvon, S.D., and Sander, J.W. (2003). A prospective study o0f the requirement for and the provision of epilepsy surgery in the United Kingdom. Epilepsia 44, 673–676.

Lin, E.-J.D., Young, D., Baer, K., Herzog, H., and During, M.J. (2006). Differential actions of NPY on seizure modulation via Y and Y receptors: evidence from receptor knockout mice. Epilepsia 47, 773–780.

Lobb, C. (2014). Abnormal Bursting as a Pathophysiological Mechanism in Parkinson’s Disease. Basal Ganglia 3, 187–195.

Lundberg, C., Björklund, T., Carlsson, T., Jakobsson, J., Hantraye, P., Déglon, N., and Kirik, D. (2008). Applications of lentiviral vectors for biology and gene therapy of neurological disorders. Curr. Gene Ther. 8, 461–473.

McCown, T.J. (2006). Adeno-associated Virus-Mediated Expression and Constitutive Secretion of Galanin Suppresses Limbic Seizure Activity in Vivo. Mol Ther 14, 63–68.

Ngugi, A.K., Bottomley, C., Kleinschmidt, I., Sander, J.W., and Newton, C.R. (2010). Estimation of the burden of activeand life-time epilepsy: a meta-analytic approach. Epilepsia 51, 883–890.

Nikitidou, L., Torp, M., Fjord-Larsen, L., Kusk, P., Wahlberg, L.U., and Kokaia, M. (2014). Encapsulated galanin-producing cells attenuate focal epileptic seizures in the hippocampus. Epilepsia 55, 167–174.

Noé, F., Pool, A.-H., Nissinen, J., Gobbi, M., Bland, R., Rizzi, M., Balducci, C., Ferraguti, F., Sperk, G., During, M.J., et al. (2008). Neuropeptide Y gene therapy decreases chronic spontaneous seizures in a rat model of temporal lobe epilepsy. Brain 131, 1506–1515.

Palfi, S., Gurruchaga, J.M., Ralph, G.S., Lepetit, H., Lavisse, S., Buttery, P.C., Watts, C., Miskin, J., Kelleher, M., Deeley, S., et al. (2014). Long-term safety and tolerability of ProSavin, a lentiviral vector-based gene therapy for Parkinson’s disease: a dose escalation, open-label, phase 1/2 trial. Lancet Lond. Engl. 383, 1138–1146.

Picot, M.-C., Baldy-Moulinier, M., Daurès, J.-P., Dujols, P., and Crespel, A. (2008). The prevalence of epilepsy and pharmacoresista0nt epilepsy in adults: a population-based study in a Western European country. Epilepsia 49, 1230–1238.

Rahim, A.A., Wong, A.M.S., Howe, S.J., Buckley, S.M.K., Acosta-Saltos, A.D., Elston, K.E., Ward, N.J., Philpott, N.J., Cooper, J.D., Anderson, P.N., et al. (2009). Efficient gene delivery to the adult and fetal CNS usingpseudotyped non-integrating lentiviral vectors. Gene Ther. 16, 509–520.

Richichi, C., Lin, E.-J.D., Stefanin, D., Colella, D., Ravizza, T., Grignaschi, G., Veglianese, P., Sperk, G., During, M.J., and Vezzani, A. (2004). Anticonvulsant and antiepileptogenic effects mediated by adeno-associated virus vector neuropeptide Y expression in the rat hippocampus. J. Neurosci. Off. J. Soc. Neurosci. 24, 3051–3059.

Schuele, S.U., and Lüders, H.O. (2008). Intractable epilepsy: management and therapeutic alternatives. Lancet Neurol. 7, 514–524.

Snowball, A., and Schorge, S. (2015). Changing channels in pain and epilepsy: Exploiting ion channel gene therapyfor disorders of neuronal hyperexcitability. FEBS Lett. 589, 1620–1634.

Suwanmanee, T., Hu, G., Gui, T., Bartholomae, C.C., Kutschera, I., von Kalle, C., Schmidt, M., Monahan, P.E., and Kafri, T. (2014). Integration-deficient lentiviral vectors expressing codon-optimized R3L human FIX restore normal hemostasis in Hemophilia B mice. Mol. Ther. J. Am. Soc. Gene Ther. 22, 567–574.

Tomlinson, S.E., Rajakulendran, S., Tan, S.V., Graves, T.D., Bamiou, D.E., Labrum, R.W., Burke, D., Sue, C.M., Giunti, P., Schorge, S., et al. (2013). Clinical, genetic, neurophysiological and functional study of new mutations in episodic ataxia type 1. J Neurol Neurosurg Psychiatry.

Woldbye, D.P.D., Angehagen, M., GØtzsche, C.R., ElbrØnd-Bek, H., SØrensen, A.T., Christiansen, S.H., Olesen, M.V., Nikitidou, L., Hansen, T.V.O., Kanter-Schlifke, I., et al. (2010). Adeno-associated viral vector-induced overexpression of neuropeptide Y Y receptors in the hippocampus suppresses seizures. Brain J. Neurol. 133, 2778–2788.

Wykes, R.C., Heeroma, J.H., Mantoan, L., Zheng, K., MacDonald, D.C., Deisseroth, K., Hashemi, K.S., Walker, M.C., Schorge, S., and Kullmann, D.M. (2012). Optogenetic and potassium channel gene therapy in arodent model of focal neocortical epilepsy. Sci. Transl. Med. 4, ra152.

Yaguchi, M., Ohashi, Y., Tsubota, T., Sato, A., Koyano, K.W., Wang, N., and Miyashita, Y. (2013). Characterization of the properties of seven promoters in the motor cortex of rats and monkeys after lentiviralvector-mediated gene transfer. Hum. Gene Ther. Methods 24, 333–344.

Yáñez-Muñoz, R.J., Balaggan, K.S., MacNeil, A., Howe, S.J., Schmidt, M., Smith, A.J., Buch, P., MacLaren, R.E., Anderson, P.N., Barker, S.E., et al. (2006). Effective gene therapy with nonintegrating lentiviral vectors. Nat. Med. 12, 348–353.

